# Microstructural and cellular characterisation of the subchondral trabecular bone in human knee and hip osteoarthritis using synchrotron tomography

**DOI:** 10.1101/2023.01.18.524641

**Authors:** Dzenita Muratovic, David M. Findlay, Micaela J. Quinn, Ryan D. Quarrington, Lucian B. Solomon, Gerald J. Atkins

## Abstract

**Objective:** It is unclear if different factors influence osteoarthritis (OA) progression and the changes characterising OA disease in hip and knee. We investigated the difference between hip OA and knee OA at the subchondral bone tissue and cellular level, relative to the degree of cartilage degeneration.

**Design:** Bone samples were collected from 11 patients (aged 70±8 years) undergoing knee arthroplasty and 8 patients (aged 64±12 years) undergoing hip arthroplasty surgery. Bone microstructure, osteocyte-lacunar network and bone matrix vascularity were evaluated using synchrotron micro-CT imaging. Samples were additionally examined histologically to determine osteocyte density, viability, and connectivity.

**Results:** After adjustment for donor gender and age, associations between the extent of cartilage degeneration, bone volume fraction [8.7, 95% CI (3.4, 14.1)], trabecular number [1.5, 95% CI (0.8, 2.3)], osteocyte lacunar density [4714.9; 95% CI (2079.1, 7350.6)] and trabecular separation [-0.06, 95% CI (0.01, 0.1)] were found in both knee and hip OA.

When compared to knee OA, hip OA was characterised by higher trabecular thickness [0.006, 95% CI (-4, 0.01)], larger but less spheric osteocyte lacunae [47.3; 95% CI (11.2, 83.4), -0.04; 95% CI (-0.6, -0.01), respectively], lower vascular canal density [-22.8; 95% CI (-35.4, -10.3)] lower osteocyte density [-84.9; 95% CI (-102.4, -67.4)], and less senescent but more apoptotic osteocytes [-2.4; 95% CI (-3.6, -1.2), 24.9; 95% CI (17.7, 32.1)], respectively.

**Conclusion:** Subchondral bone from hip OA and knee OA exhibits different characteristics at the tissue and cellular levels, suggesting different mechanisms of OA progression between the hip and knee joints.

## Introduction

Osteoarthritis (OA) is a multifactorial joint disease that is most frequently seen in the knee, hip, hand, and spine ^1, 2^. It is unclear whether the aetiology of OA is joint-specific, or if the risk factors that influence OA progression and structural degeneration are common between joints.

Hip and knee osteoarthritis are the most common forms of OA. The clinical presentation of patients, including age, gender, symptom duration, pain medication usage, pain intensity, physical function and quality of life, are alike for hip and knee OA ^3^. While the prevalence of hip OA is lower than knee OA, it has been reported that hip replacements are performed in younger, predominantly male individuals, with lower mean BMI than the knee OA population ^4^. Although this may be explained by the relative success of total hip replacement (THR) over total knee replacement (TKR) in this population, it is also possible that OA in the hip progresses more rapidly than in the knee.

Studies have also indicated different gene and molecular marker expression in hip and knee OA. For example, homeobox (HOX) genes and certain micro-RNAs (miRNAs) were found to be differentially expressed in articular cartilage between hip and knee OA, ^5, 6^ as well as inflammatory cytokine and bone metabolism markers in serum and synovial fluid ^7, 8^. More recently, Styrkarsdottir et al., in the largest genome-wide association study (GWAS) for OA disease including 17,151 hip osteoarthritis patients, 23,877 knee osteoarthritis patients and 562,000 controls, identified 16 new loci: 12 for hip and 4 for knee osteoarthritis, without overlap. This further supported the hypothesis that hip and knee OA are different OA subtypes with different genetic drivers ^9^.

Importantly, while there are many studies investigating the association between structural degeneration, clinical symptoms and imaging findings for the subchondral bone (SCB) in knee OA, that of hip OA is relatively understudied ^10^. Currently, there are no studies directly comparing the structural changes in osteochondral tissue between the knee and hip in the context of OA. Therefore, little is known about the comparative underlying mechanisms that drive OA progression in these joints.

This study was designed to investigate the type of structural SCB changes between hip and knee OA at the tissue level, using high-resolution synchrotron tomography. This imaging modality provides quantitative, in-depth information of bone microarchitecture and bone microporosity, consisting largely of vascular channels and the lacunar network occupied by the predominant bone cell type, the osteocyte. This analysis was complemented by a histological assessment of osteocyte viability, apoptosis and senescence in the same specimens.

## Materials and Methods

### Clinical Specimens

Tibial plateaus were collected from 11 patients (6 males, 66±9; 5 females, 70±8 years) undergoing TKR surgery and 8 femoral heads (4 males, 69±14; 4 females, 64±12 years) from patients who had undergone THR surgery. Inclusion criteria were end-stage OA based on radiographic imaging and symptomatic disabilities. Exclusion criteria were trauma, rheumatoid arthritis, osteoporosis, malignancy and metabolic bone disease. Written consent was obtained from all participants and the study received prior approval from the Human Research Ethics Committee at the Royal Adelaide Hospital and The University of Adelaide (Approval No. HS-2013-003). Upon receiving surgical samples, two bone core biopsies (8 × 7 mm) were taken from each sample. For knee samples, the biopsies were obtained from the anterior and posterior aspects of the medial condyles of the tibial plateaus (Fig.1A). From hip samples, the biopsies were obtained from the anterior-superior and anterior-inferior aspects of the femoral head (Fig 1B).

**Figure 1.**
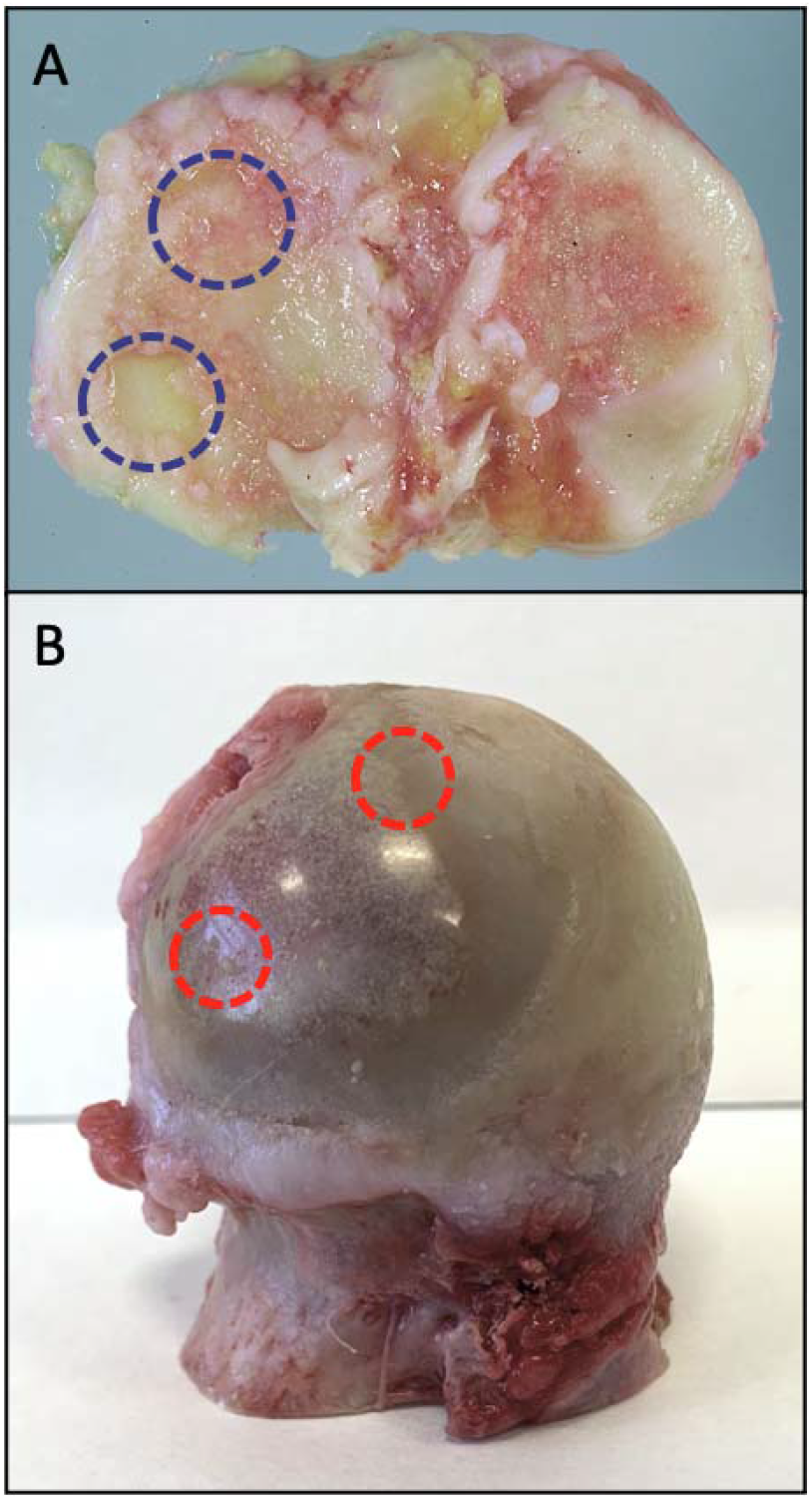
Gross view of an osteoarthritic tibial plateau, female, age 76 years (A) and femoral head, female, age 75 years (B) from patients who had undergone total joint replacement surgery. Red and blue circles indicate biopsy sites.

### Synchrotron radiation micro-computed tomography (SRμCT)

Synchrotron tomographic X-ray experiments were carried out using the X02DA TOMCAT beamline of the Swiss Light Source (SLS) facility at the Paul Scherrer Institute. Before imaging, bone biopsies were fixed in formaldehyde for 24h, rinsed several times in 75% ethanol and embedded in paraffin to be protected from overheating damage during scanning. Bone cores were scanned with a monochromatic beam of 20 keV. The X-ray imaging system consisted of a microscope (Optique Peter) with a 4x magnification objective and LuAG: Ce 20μm scintillator coupled to a pco.edge 5.5 detector with a pixel size of 1.63 μm and a field of view 4.2×3.5 mm^2^. For each sample, 1800 projections, uniformly distributed over 180 degrees, were taken with 120 ms exposure each, resulting in total scan times of approximately 6 minutes. Tomographic reconstructions were performed on the TOMCAT cluster ^11^ using a phase retrieval method from a single defaced image ^12^ and the GridRec algorithm ^13^. Final reconstructed images were saved as 16-bit TIFF image stacks.

Image stacks were analysed using CTAnalyser (v.1.18.8.0, Bruker, Kontich, Belgium). Cylindrical regions of interest (ROI) measuring 3 mm in diameter and 1 mm in height and an approximate volume of 7 mm^3^, of trabecular bone, were analysed to obtain trabecular bone microstructure (Fig. 2), density, size (Fig. 3A & 3B), and shape of osteocyte lacunae and density of vascular canals in bone matrix (Fig. 3C & 3D). The following trabecular bone microstructure parameters were assessed: bone volume fraction (BV/TV), Trabecular Number (Tb. N), Trabecular Thickness (Tb. Th) and Trabecular Separation (Tb. Sp). Bone mineral density was evaluated from the original reconstructed images, using a custom-written MATLAB code, as previously described ^14^.

**Figure 2.**
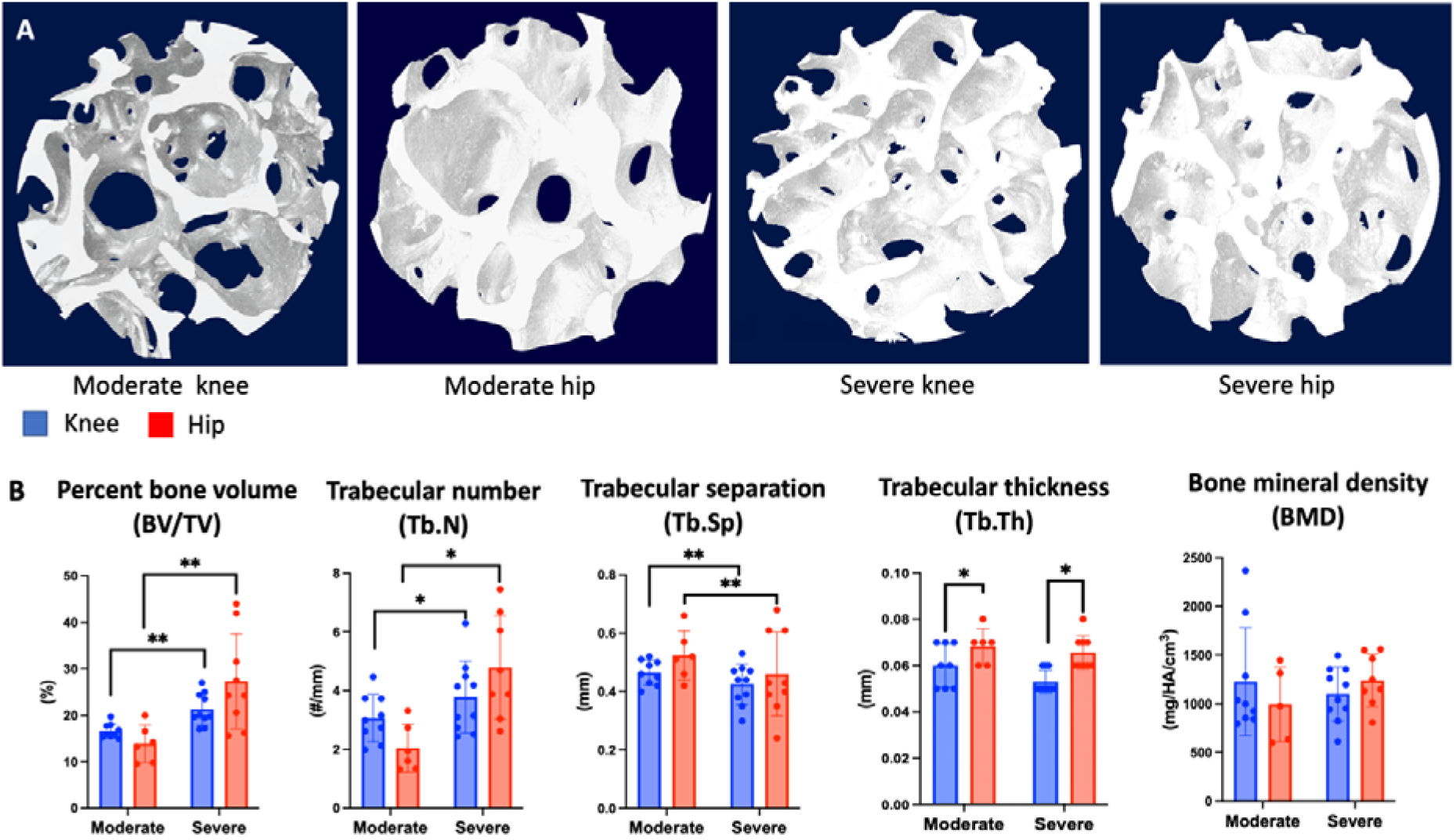
A) Reconstructed 3D models from Synchrotron Radiation X-ray micro-computed tomography images of subchondral bone microstructure in the moderate and severe knee and hip OA. B) Quantified microarchitectural measures. Each point represents an individual sample. Significant differences are indicated by *p<0.05, **p<0.005.

**Figure 3.**
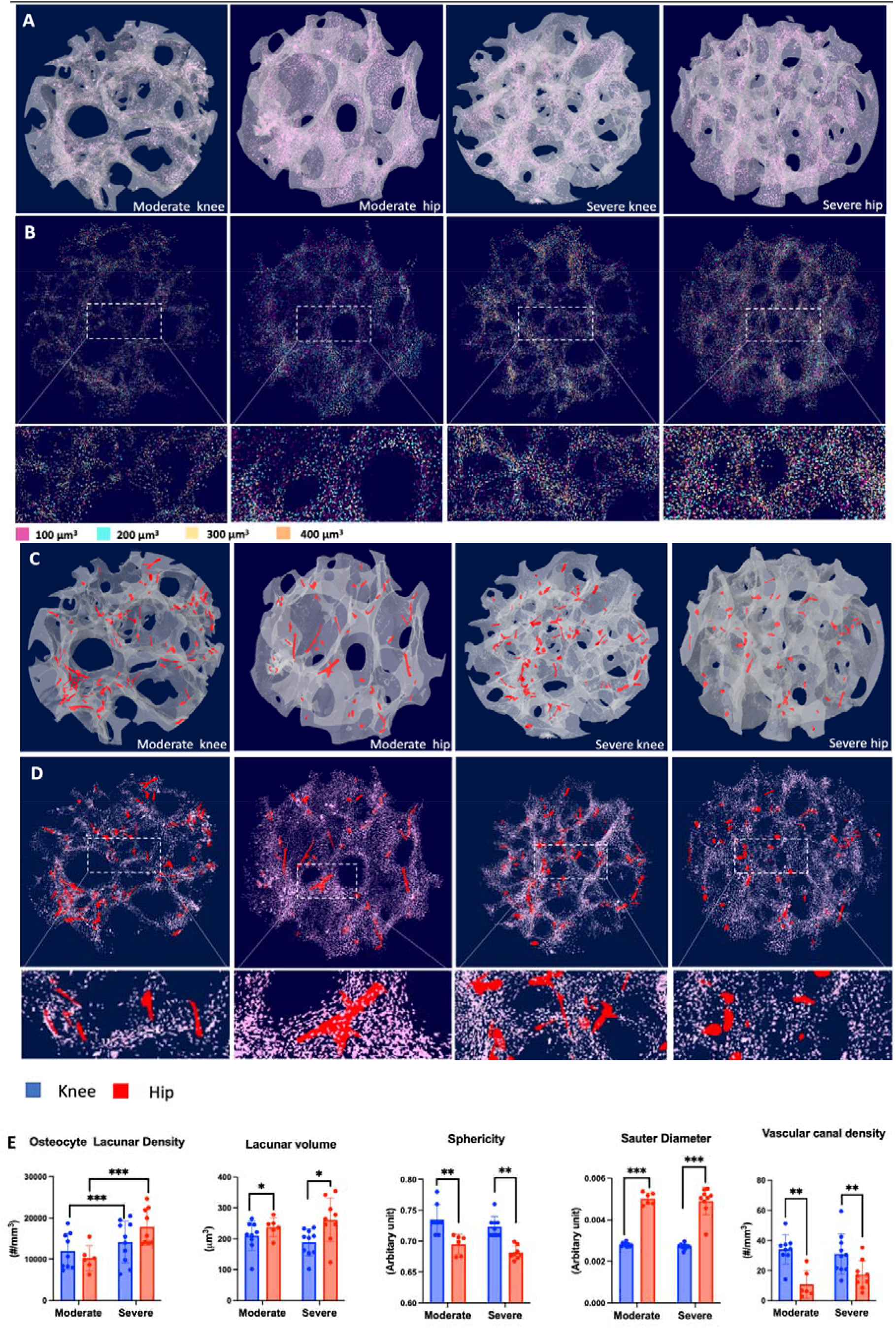
Reconstructed 3D models from Synchrotron Radiation X-ray micro-computed tomography images A) the density of osteocyte lacunae (pink), B) colour-coded size of lacunae and C&D) vascular canals (red) within the bone matrix. E) Quantified osteocyte lacunae measures. Each point represents an individual sample. Significance is indicated by *p<0.05, **p<0.005, ***p<0.0005.

Osteocyte lacunae and vascular canals were identified using 2-step Otsu thresholding. Osteocyte lacunae were defined as closed, pore-like structures above 30 μm^3^ and less than 2000 μm^3^. Osteocyte lacunar parameters lacunar density (La.Dn, #/mm^2^), lacunar volume (La.V, μm^3^), orientation of lacunae (°), Sauter diameter (SD) and Sphericity (Sph) were obtained using individual object analysis. Vascular canals within the bone matrix were defined as open pore-like structures greater than 2000 μm^3^. The following parameters for the vascular canal were determined: canal density (canal number/bone volume, #/mm^3^), total canal volume (*μ*m^3^) and mean vascular canal diameter (*μ*m).

### Histology

Following synchrotron imaging, bone core samples were slow-decalcified in 10% ethylenediaminetetraacetic acid (EDTA) solution, processed, and embedded in paraffin. 20 sections were cut per sample (5*μ*m thickness) and prepared for histological staining.

Degenerative changes in the osteochondral unit (cartilage and bone) were evaluated using the Osteoarthritis Research Society International (OARSI) grading system, on tissue sections stained with Safranin O Fast Green. Osteocyte cell density was quantified on sections stained with Haematoxylin and Eosin (Fig. 4A). For visualisation of the osteocyte lacunocanalicular network and to evaluate average canalicular number and length, Ploton silver staining ^15^ was used (Fig. 4B). According to previously published protocols ^16, 17^, Sudan Black B staining was used for visualisation and quantitation of senescent osteocyte density (Fig. 4C). Osteocytes undergoing apoptosis were detected in terminal deoxynucleotidyl transferase dUTP nick end labelling (TUNEL) assays ^18^, using a commercially available In Situ Cell Death Detection kit (Sigma-Aldrich), according to the manufacturer’s protocol (Fig. 4D). Each analysis was performed in two separate sections and in 5 ROI per section. Reported parameters were then determined by calculating average value per mm^2^ of bone tissue.

**Figure 4.**
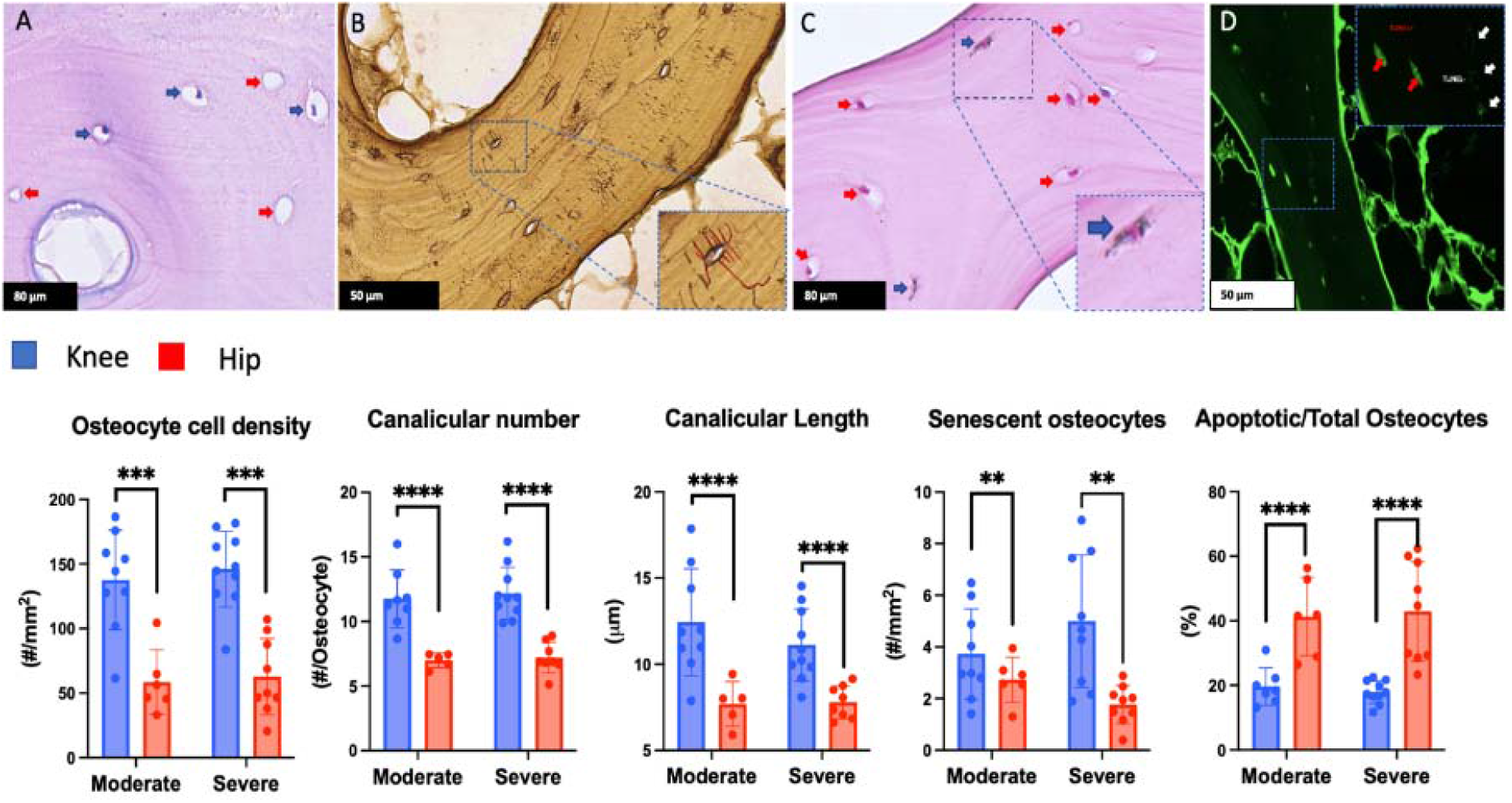
Histological assessment of osteocyte viability, senescence and connectivity. (A) H&E staining, (B) Ploton silver staining, (C) Sudan Black B staining (D) terminal deoxynucleotidyl transferase dUTP nick end labelling (TUNEL) assays. Each point represents an individual sample. Significance is indicated by **p<0.005, ***p<0.0005, ****p<0.0001.

### Statistical analysis

Statistics were performed using SPSS v28 (IBM, Illinois, USA). Separate linear regression models with cluster-robust standard errors assessed *a priori* whether each structural and cellular bone parameter differed due to joint type (knee or hip) and OARSI grade (moderate vs severe), or the interaction of the two, when adjusted for donor gender and age. For each model, if the joint*OARSI interaction was not significantly associated with the outcome then it was removed so that the effect of each *a priori* variable was assessed independently. Cluster-robust standard errors accounted for non-independence due to multiple samples being analysed from the same donor tissue.

## Results

### Bone Microstructure

After adjustment for donor gender and age, positive associations between OARSI grade and bone volume fraction [estimate=8.7, 95% CI (3.4, 14.1), p<0.005], trabecular number [estimate=1.5, 95% CI (0.8, 2.3), p<0.001] and decreased trabecular separation [estimate=-0.06, 95% CI (0.01, 0.1), p<0.005] were found in both hip and knee OA.

The hip OA had a significantly higher trabecular thickness [estimate=0.006, 95% CI (−4, 0.01), p<0.05] compared to knee OA independently of OARSI grade. No significant interaction was detected between OARSI grade and type of joint with bone mineral density (Fig. 2B).

### Osteocyte Lacunar density and morphometric characteristics

Disease severity had a significant association with increased osteocyte lacunar density [estimate=4714.9; 95% CI (2079.1, 7350.6) p<0.001], independently of type of joint and patient age. The type of joint had significant associations with lacunar volume [estimate=47.3; 95% CI (11.2, 83.4), p<0.05], sphericity [estimate=-0.04; 95% CI (−0.6, -0.01), p<0.005], and Sauter diameter [estimate=0.002; 95% CI (0.002, 0.002), p<0.0001] independently of OARSI grade and patient age. Specifically, larger lacunae that are flatter and less spherical were found in SCB of hip OA when compared to that of knee OA. After adjustment for patient age and gender, lacunar orientation was not significantly impacted by OARSI grade, or with type of the joint. Female gender was positively associated with osteocyte lacunar density [estimate=4954.1; 95% CI (1873.7, 8034.4) p<0.005] and with lacunar volume [estimate=39.7; 95% CI (6.9, 72.4), p<0.05], (Fig 3E).

### Bone Matrix Vascularity

Trabecular bone in hip OA samples had lower vascular canal density [-22.8; 95% CI (−35.4, -10.3)] compared to knee OA, independent of disease severity and gender (Fig 3E). Further, a negative association was found between increasing patient age and vascular canal density [estimate=-0.7, 95% CI (−1.1, -0.2), p<0.005]. No statistically significant interaction was detected between OARSI grade and type of joint with total canal volume and mean vascular canal diameter.

### Osteocyte cell viability and connectivity

The type of joint had a significant effect on cellular parameters in SCB independent of OARSI grade and patient age. Specifically, lower osteocyte density [estimate=-84.9; 95% CI (−102.4, -67.4), p<0.0005], lower density of senescent osteocytes [estimate=-2.4; 95% CI (−3.6, -1.2), p<0.005] and higher percentage of apoptotic cells [estimate=24.9; 95% CI (17.7, 32.1), p<0.005] and percentage of empty lacunae [estimate=39.1; 95% CI (32.9, 45.3), p<0.0001] were found in the SCB of hip OA when compared to knee OA.

Consistent with the increased density of apoptotic osteocytes, lower average canalicular number per osteocyte [estimate=-4.7; 95% CI (−5.7, -3.7), p<0.0001] and canalicular length [estimate=-4; 95% CI (−5.2, -2.8), p<0.0001], indicating decreased connectivity between osteocytes, were also found in SCB of hip OA independent of OARSI grade, patient age and gender.

## Discussion

In this study, we measured micro-structural and cellular parameters of SCB tissue from individuals with either hip or knee OA, comparing bone from zones of moderate and severe OA, as defined by the quality (OARSI grade) of the overlying cartilage. Our results indicate that while bone volume fraction, trabecular number and trabecular separation were altered according to disease severity in both joints, hip and knee SCB exhibited distinct characteristics at the osteocyte lacunocanalicular level, independently of disease severity, suggesting different mechanisms of disease progression.

Physiologic mechanical loading is required for optimal joint health, including cartilage renewal ^19^ and SCB remodelling ^20^. It is well accepted that mechanical loading has an essential role in bone metabolism and structural adaptation, and that over- or aberrant loading plays a key role in the onset and OA progression. SCB sclerosis along with progressive cartilage degradation, is a hallmark of late stage OA ^21^ and is considered to be a direct adaptive response of bone to persistent mechanical over-loading. Consistent with this, we found in both hip and knee OA a sclerotic appearance of bone in association with increased OARSI grade. Further analysis showed that an increased OARSI grade had a statistically significant effect on the sclerotic appearance of SCB (increase of percent bone volume and trabecular number and decrease of trabecular separation) in both hip and knee OA, while the skeletal site has a statistically significant effect on increased trabecular thickness. This suggests that the adaptation of bone microstructure may occur differently in hip and knee OA. A possible explanation for this phenomenon is that the hip and knee are different types of synovial joint (ball and socket vs hinge), with different biomechanics ^4, 22^ and likely, therefore, different aetiologies of disease progression.

Osteocytes have been identified as the cells primarily responsible for sensing mechanical loading of bone ^23^ and in response orchestrate the activity of osteoblast or osteoclasts to remodel or model bone ^23^. Osteocytes also maintain the health of the bone by regulating bone matrix mineralisation ^24, 25^.

In this study, SCB differences were seen between hip and knee OA in terms of osteocyte cell density, viability, senescence, and the lacunocanalicular network. These results may indicate underlying differences in osteocyte behaviour in the two skeletal sites, or perhaps a compromised ability to respond to the environment imposed by the OA disease state. Previously, an increase in mean osteocyte lacunar density but with decreased osteocyte viability in knee OA was reported ^18, 26^ and in human hip OA, increased numbers of immature osteocytes were associated with sclerotic bone ^27^. In addition, a decline in osteocyte viability has been associated with increasing age and the accumulation of microcracks within bone tissue of humans ^28, 29^ and mice ^30^.

The accumulation of microdamage, including microcracks, diffuse damage and micro-fractures in trabecular bone is closely associated with mechanical over-loading (amount and frequency). Previously, we reported evidence of increased accumulation of linear microcracks and diffuse damage in knee OA ^26^, while Taljanovic et al. reported the presence of micro-fractures in hip OA ^31^. The repair of microfractures in the damaged region of bone matrix is thought to be initiated by osteocyte apoptosis triggering osteoclastic bone resorption ^32^. In contrast, diffuse damage is repaired by an unknown osteocyte-mediated mechanism ^33^. In this study, increased osteocyte apoptosis was detected in SCB of hip OA compared to knee OA. This may suggest that SCB in the hip OA has reduced ability to repair and hence displays more severe tissue changes in overload-induced OA progression.

Using synchrotron micro-CT, we have been able to obtain high resolution, 3D morphometric characteristics of osteocyte lacunae. We found that hip OA is characterised with larger lacunae that are less spherical and more flat (higher sauter diameter) compared to knee OA.

The amount, type and direction of the loading has a direct effect on the size and shape of osteocyte lacunae ^34^. In animal models, it has been reported that osteocyte lacunar volume increases or decreases with increased or decreased loading ^34, 35^.

Further, osteocyte lacunae tend to be smaller and more spherical in bones with low impact loading compared to bones with high impact loading ^36^. In addition, it is hypothesised that changes to lacunar morphology may influence the mechanosensitivity of the osteocyte. Specifically, larger lacunae contain osteocytes with increased mechanosensitivity ^37^.

In addition to 3D lacunar morphometry, we also demonstrated a significantly lower vascular canal density within the SCB in hip OA compared to knee OA. This is of interest because osteonecrosis was proposed to be a direct cause of hip OA ^38^. Moreover, it was suggested that osteonecrosis may occur in the SCB many years before clinical presentation of OA ^38-40^. More recent studies have also reported the close relationship between OA and osteonecrosis ^41^.

A role for vascular pathology in the development and progression of OA was suggested by Findlay ^42^, proposing the potential value for further investigation of vascular-directed treatments to inhibit the progression of OA. More recently, clinical studies have demonstrated a causal relationship between vascular pathology and knee and hand OA but there are no data for hip OA ^43^.

Taken together, findings of reduced osteocyte cell density and/or a higher ratio of empty over total lacunae (74%), increased apoptosis and decreased vascularity, compared with knee OA, are supportive that tissue necrosis in the SCB in hip OA has important clinical implications.

Ageing is a common risk factor in the development and progression of human knee and hip OA. Animal studies have shown that age is a major factor of influencing the ability of bone to heal and regain mechanical competence after fracture ^44^. Age reduces the rate of bone tissue cellular repair and differentiation and the rate of revascularisation ^45, 46 47^. One of the hallmarks of ageing is the accumulation of the senescent cells, which can adversely affect tissue healing. Recent studies provide evidence of senescent cells in cartilage during OA development ^48-50^, while the role of SCB cell senescence in OA progression has not yet been described. In this study, we used a specific histochemical stain for Lipofuscin, a tissue biomarker indicative of stress-induced senescence ^16 17^. Interestingly, we found that the SCB of knee OA contained lipofuscin-positive/senescent cells to a greater extent than SCB of hip OA, independent of the degree of disease progression. It is possible that an increased number of senescent osteocytes is a feature of the knee SCB irrespective of OA, however this accumulation might nevertheless affect tissue repair in OA progression in the knee. However, since the presence of lipofuscin is not an absolute marker of senescence, further investigation is needed.

We believe that this is the first study to compare directly the tissue characteristics of the SCB in hip and knee OA. However, this study has several limitations. Firstly, the number of samples is relatively small. Secondly, the absence of control tissue for the hip and our inability to include early OA disease samples from either joint, limits our conclusions.

In summary, we observed significant differences in osteocyte cellularity and connectivity, and vascularity in hip OA SCB, compared to SCB from knee OA. The results suggest that while OA of the hip and knee is characterised by similar changes to bone microarchitecture, with the exception of trabecular thickness, potentially different disease mechanisms are responsible. An implication of these findings is that specific approaches to treat disease progression may be needed for OA in different joints.

## Acknowledgements

The authors acknowledge the Paul Scherrer Institut, Villigen, Switzerland for provision of the synchrotron radiation beam time at the TOMCAT beamline of the Swiss Light Source and would like to particularly thank to Dr. Elena Borisova and Dr Christian Schlepuetz for their support during and after beamtime at TOMCAT.

We also acknowledge travel funding provided by the International Synchrotron Access Program (ISAP) managed by the Australian Synchrotron, part of ANSTO, and funded by the Australian Government.

The authors wish to acknowledge support from Adelaide Microscopy at The University of Adelaide, especially Dr Agatha Labrinidis, for assisting with developing the method to obtain data from Synchrotron micro-CT images.

## Author contributions

All authors meet the criteria for authorship. DM designed the study, performed all experiments and analysed the results, interpreted the data, and drafted the manuscript. MQ performed experiments and analysis of the results, RQ performed statistical analysis and developed MATLAB code/programs for data collection and interpreted the data. DF, BS and GA interpreted the data and provided overall supervision. All authors contributed to manuscript editing and approved submitted version.

## References

1. Bortoluzzi A, Furini F, Scire CA. Osteoarthritis and its management - Epidemiology, nutritional aspects and environmental factors. Autoimmun Rev 2018; 17: 1097–1104.

2. Primorac D, Molnar V, Rod E, Jelec Z, Cukelj F, Matisic V, et al. Knee Osteoarthritis: A Review of Pathogenesis and State-Of-The-Art Non-Operative Therapeutic Considerations. Genes (Basel) 2020; 11.

3. Roos EM, Gronne DT, Thorlund JB, Skou ST. Knee and hip osteoarthritis are more alike than different in baseline characteristics and outcomes: a longitudinal study of 32,599 patients participating in supervised education and exercise therapy. Osteoarthritis Cartilage 2022; 30: 681–688.

4. Bennell KL, Hunt MA, Wrigley TV, Hunter DJ, McManus FJ, Hodges PW, et al. Hip strengthening reduces symptoms but not knee load in people with medial knee osteoarthritis and varus malalignment: a randomised controlled trial. Osteoarthritis Cartilage 2010; 18: 621–628.

5. Coutinho de Almeida R, Ramos YFM, Mahfouz A, den Hollander W, Lakenberg N, Houtman E, et al. RNA sequencing data integration reveals an miRNA interactome of osteoarthritis cartilage. Ann Rheum Dis 2019; 78: 270–277.

6. Le Douarin NM, Creuzet S, Couly G, Dupin E. Neural crest cell plasticity and its limits. Development 2004; 131: 4637–4650.

7. Ren G, Lutz I, Railton P, Wiley JP, McAllister J, Powell J, et al. Serum and synovial fluid cytokine profiling in hip osteoarthritis: distinct from knee osteoarthritis and correlated with pain. BMC Musculoskelet Disord 2018; 19: 39.

8. Van Spil WE, Welsing PM, Bierma-Zeinstra SM, Bijlsma JW, Roorda LD, Cats HA, et al. The ability of systemic biochemical markers to reflect presence, incidence, and progression of early-stage radiographic knee and hip osteoarthritis: data from CHECK. Osteoarthritis Cartilage 2015; 23: 1388–1397.

9. Styrkarsdottir U, Lund SH, Thorleifsson G, Zink F, Stefansson OA, Sigurdsson JK, et al. Meta-analysis of Icelandic and UK data sets identifies missense variants in SMO, IL11, COL11A1 and 13 more new loci associated with osteoarthritis. Nat Genet 2018; 50: 1681–1687.

10. Hall M, van der Esch M, Hinman RS, Peat G, de Zwart A, Quicke JG, et al. How does hip osteoarthritis differ from knee osteoarthritis? Osteoarthritis Cartilage 2022; 30: 32–41.

11. Hintermuller C, Marone F, Isenegger A, Stampanoni M. Image processing pipeline for synchrotron-radiation-based tomographic microscopy. J Synchrotron Radiat 2010; 17: 550–559.

12. Paganin D, Mayo SC, Gureyev TE, Miller PR, Wilkins SW. Simultaneous phase and amplitude extraction from a single defocused image of a homogeneous object. J Microsc 2002; 206: 33–40.

13. Marone F, Stampanoni M. Regridding reconstruction algorithm for real-time tomographic imaging. J Synchrotron Radiat 2012; 19: 1029–1037.

14. Muratovic D, Findlay DM, Quarrington RD, Cao X, Solomon LB, Atkins GJ, et al. Elevated levels of active Transforming Growth Factor beta1 in the subchondral bone relate spatially to cartilage loss and impaired bone quality in human knee osteoarthritis. Osteoarthritis Cartilage 2022; 30: 896–907.

15. Dole NS, Yee CS, Schurman CA, Dallas SL, Alliston T. Assessment of Osteocytes: Techniques for Studying Morphological and Molecular Changes Associated with Perilacunar/Canalicular Remodeling of the Bone Matrix. Methods Mol Biol 2021; 2230: 303–323.

16. Evangelou K, Gorgoulis VG. Sudan Black B, The Specific Histochemical Stain for Lipofuscin: A Novel Method to Detect Senescent Cells. Methods Mol Biol 2017; 1534: 111–119.

17. Georgakopoulou EA, Tsimaratou K, Evangelou K, Fernandez Marcos PJ, Zoumpourlis V, Trougakos IP, et al. Specific lipofuscin staining as a novel biomarker to detect replicative and stress-induced senescence. A method applicable in cryo-preserved and archival tissues. Aging (Albany NY) 2013; 5: 37–50.

18. Jaiprakash A, Prasadam I, Feng JQ, Liu Y, Crawford R, Xiao Y. Phenotypic characterization of osteoarthritic osteocytes from the sclerotic zones: a possible pathological role in subchondral bone sclerosis. Int J Biol Sci 2012; 8: 406–417.

19. Lim CT, Zhou EH, Quek ST. Mechanical models for living cells--a review. J Biomech 2006; 39: 195–216.

20. Findlay DM, Atkins GJ. Osteoblast-chondrocyte interactions in osteoarthritis. Curr Osteoporos Rep 2014; 12: 127–134.

21. Burr DB, Gallant MA. Bone remodelling in osteoarthritis. Nat Rev Rheumatol 2012; 8: 665–673.

22. Reijman M, Pols HA, Bergink AP, Hazes JM, Belo JN, Lievense AM, et al. Body mass index associated with onset and progression of osteoarthritis of the knee but not of the hip: the Rotterdam Study. Ann Rheum Dis 2007; 66: 158–162.

23. Bonewald LF. The amazing osteocyte. J Bone Miner Res 2011; 26: 229–238.

24. Robling AG, Bonewald LF. The Osteocyte: New Insights. Annu Rev Physiol 2020; 82: 485–506.

25. Prideaux M, Findlay DM, Atkins GJ. Osteocytes: The master cells in bone remodelling. Curr Opin Pharmacol 2016; 28: 24–30.

26. Muratovic D, Findlay DM, Cicuttini FM, Wluka AE, Lee YR, Kuliwaba JS. Bone matrix microdamage and vascular changes characterize bone marrow lesions in the subchondral bone of knee osteoarthritis. Bone 2018; 108: 193–201.

27. Ilas DC, Churchman SM, Baboolal T, Giannoudis PV, Aderinto J, McGonagle D, et al. The simultaneous analysis of mesenchymal stem cells and early osteocytes accumulation in osteoarthritic femoral head sclerotic bone. Rheumatology (Oxford) 2019; 58: 1777–1783.

28. Vashishth D, Verborgt O, Divine G, Schaffler MB, Fyhrie DP. Decline in osteocyte lacunar density in human cortical bone is associated with accumulation of microcracks with age. Bone 2000; 26: 375–380.

29. Busse B, Djonic D, Milovanovic P, Hahn M, Puschel K, Ritchie RO, et al. Decrease in the osteocyte lacunar density accompanied by hypermineralized lacunar occlusion reveals failure and delay of remodeling in aged human bone. Aging Cell 2010; 9: 1065–1075.

30. Tiede-Lewis LM, Xie Y, Hulbert MA, Campos R, Dallas MR, Dusevich V, et al. Degeneration of the osteocyte network in the C57BL/6 mouse model of aging. Aging (Albany NY) 2017; 9: 2190–2208.

31. Taljanovic MS, Graham AR, Benjamin JB, Gmitro AF, Krupinski EA, Schwartz SA, et al. Bone marrow edema pattern in advanced hip osteoarthritis: quantitative assessment with magnetic resonance imaging and correlation with clinical examination, radiographic findings, and histopathology. Skeletal Radiol 2008; 37: 423–431.

32. Kennedy OD, Laudier DM, Majeska RJ, Sun HB, Schaffler MB. Osteocyte apoptosis is required for production of osteoclastogenic signals following bone fatigue in vivo. Bone 2014; 64: 132–137.

33. Herman BC, Cardoso L, Majeska RJ, Jepsen KJ, Schaffler MB. Activation of bone remodeling after fatigue: differential response to linear microcracks and diffuse damage. Bone 2010; 47: 766–772.

34. Bozal CB, Sanchez LM, Mandalunis PM, Ubios AM. Histomorphometric study and three-dimensional reconstruction of the osteocyte lacuno-canalicular network one hour after applying tensile and compressive forces. Cells Tissues Organs 2013; 197: 474–483.

35. Sugawara Y, Kamioka H, Ishihara Y, Fujisawa N, Kawanabe N, Yamashiro T. The early mouse 3D osteocyte network in the presence and absence of mechanical loading. Bone 2013; 52: 189–196.

36. Vatsa A, Breuls RG, Semeins CM, Salmon PL, Smit TH, Klein-Nulend J. Osteocyte morphology in fibula and calvaria --- is there a role for mechanosensing? Bone 2008; 43: 452–458.

37. Hemmatian H, Jalali R, Semeins CM, Hogervorst JMA, van Lenthe GH, Klein-Nulend J, et al. Mechanical Loading Differentially Affects Osteocytes in Fibulae from Lactating Mice Compared to Osteocytes in Virgin Mice: Possible Role for Lacuna Size. Calcif Tissue Int 2018; 103: 675–685.

38. Wong SY, Evans RA, Needs C, Dunstan CR, Hills E, Garvan J. The pathogenesis of osteoarthritis of the hip. Evidence for primary osteocyte death. Clin Orthop Relat Res 1987: 305–312.

39. Ghosh P, Cheras PA. Vascular mechanisms in osteoarthritis. Best Pract Res Clin Rheumatol 2001; 15: 693–709.

40. Franchi A, Bullough PG. Secondary avascular necrosis in coxarthrosis: a morphologic study. J Rheumatol 1992; 19: 1263–1268.

41. Rony L, Perrot R, Hubert L, Chappard D. Osteocyte staining with rhodamine in osteonecrosis and osteoarthritis of the femoral head. Microsc Res Tech 2019; 82: 2072–2078.

42. Findlay DM. Vascular pathology and osteoarthritis. Rheumatology (Oxford) 2007; 46: 1763–1768.

43. Hussain SM, Dawson C, Wang Y, Tonkin AM, Chou L, Wluka AE, et al. Vascular Pathology and Osteoarthritis: A Systematic Review. J Rheumatol 2020; 47: 748–760.

44. Meyer RA, Jr., Tsahakis PJ, Martin DF, Banks DM, Harrow ME, Kiebzak GM. Age and ovariectomy impair both the normalization of mechanical properties and the accretion of mineral by the fracture callus in rats. J Orthop Res 2001; 19: 428–435.

45. Lu C, Hansen E, Sapozhnikova A, Hu D, Miclau T, Marcucio RS. Effect of age on vascularization during fracture repair. J Orthop Res 2008; 26: 1384–1389.

46. Boskey AL, Coleman R. Aging and bone. J Dent Res 2010; 89: 1333–1348.

47. Boskey AL, Imbert L. Bone quality changes associated with aging and disease: a review. Ann N Y Acad Sci 2017; 1410: 93–106.

48. Gao SG, Zeng C, Li LJ, Luo W, Zhang FJ, Tian J, et al. Correlation between senescence-associated beta-galactosidase expression in articular cartilage and disease severity of patients with knee osteoarthritis. Int J Rheum Dis 2016; 19: 226–232.

49. Jeon OH, David N, Campisi J, Elisseeff JH. Senescent cells and osteoarthritis: a painful connection. J Clin Invest 2018; 128: 1229–1237.

50. Price JS, Waters JG, Darrah C, Pennington C, Edwards DR, Donell ST, et al. The role of chondrocyte senescence in osteoarthritis. Aging Cell 2002; 1: 57–65.

